# A novel metabarcoding strategy for studying nematode communities

**DOI:** 10.1101/2020.01.27.921304

**Authors:** Md. Maniruzzaman Sikder, Mette Vestergård, Rumakanta Sapkota, Tina Kyndt, Mogens Nicolaisen

## Abstract

Nematodes are widely abundant soil metazoa and often referred to as indicators of soil health. While recent advances in next-generation sequencing technologies have accelerated research in microbial ecology, the ecology of nematodes remains poorly elucidated, partly due to the lack of reliable and validated sequencing strategies. Objectives of the present study were (i) to compare commonly used primer sets and to identify the most suitable primer set for metabarcoding of nematodes; (ii) to establish and validate a high-throughput sequencing strategy for nematodes using Illumina paired-end sequencing. In this study, we tested four primer sets for amplicon sequencing: JB3/JB5 (mitochondrial, I3-M11 partition); SSU_04F/SSU_22R (18S rRNA, V1-V2 region); Nemf/18Sr2b (18S rRNA, V6-V8 region) from earlier studies; and MMSF/MMSR (18S rRNA, V4-V5 region), a newly developed primer set from this study. In order to test the primer sets, we used 22 samples of individual nematode species, 20 mock communities, 20 soil samples, 20 spiked soil samples (mock communities in soil), and 4 root/rhizosphere soil samples. We successfully amplified the target regions (I3-M11 partition of the COI gene; V1-V2, V4-V8 region of 18S rRNA gene) from these 86 DNA samples with the four different primer combinations and sequenced the amplicons on an Illumina MiSeq sequencing platform. We found that the MMSF/MMSR and Nemf/18Sr2b were efficient in detecting nematode compared to JB and SSU primer sets based on annotation of sequence reads at genus and in some cases at species level. Therefore, these primer sets are suggested for studies of nematode communities in agricultural environments.

## Background

Nematodes are highly diverse and abundant metazoans with worldwide distribution [1]. Generally, nematologists have relied on classical morphology-based taxonomy along with biochemical or molecular methods for nematode identification [2, 3]. Morphological identification is difficult, requires taxonomic expertise and often becomes challenging when it comes to identifying nematodes at lower taxonomic levels [4]. DNA based identification have eased the task of taxonomic nematode identification in recent years, and most molecular based diagnostic approaches usually target the nuclear ribosomal DNA region. In addition, the mitochondrial cytochrome oxidase I gene (COI gene) has been successfully used for identification of nematodes and for resolving taxonomic relationships among closely related species [5–7]. For certain groups of taxa, the COI gene has been shown to provide greater taxonomic resolution compared to the small subunit (SSU, 18S rRNA) rDNA [8]. The potential of COI gene-based barcoding has been explored for nematode taxa ranging from root-knot nematodes [9], marine nematodes [7], Aphelenchoididae [10] and *Pratylenchus* [11]. Both marker genes, 18S ribosomal DNA and COI, comes with their own limitations and strengths. The reference database for COI sequences is less enriched in comparison to 18S, limiting the implementation of COI barcoding for nematodes. The most inclusive molecular phylogenetic study of nematodes now available comprised 1215 full-fragment sequences of SSU rDNA [12]. There as several reports on the use of 18S rRNA based barcodes for successful nematode community analysis, and they resolved several taxonomic issues of identification of several nematodes [13–15]. Consequently, the 18S rRNA gene may remain the most widely used molecular marker for identification of nematodes [16, 17].

The field of DNA based identification is transitioning from barcoding individual species to metabarcoding of entire communities. However, the success of metabarcoding approaches largely relies on suitable primers used for amplification of environmental DNA (eDNA). Nematode community studies by earlier workers have relied on nematode extraction [18, 19] to screen out other organisms present in the samples during amplification. This process is time consuming, laborious and may introduce biases [20]. Therefore, in the present study, we compared amplification strategies that avoided such nematode isolation steps. In a previous study, we have already optimized a soil DNA extraction method that we used to evaluate nematode communities from a number of agricultural soils using the Roche 454 platform [21]. After alignment of 18S rRNA genes of eukaryotic sequences available in the SILVA database, variable regions V2, V4, and V9 were suggested as the most suitable for biodiversity assessments [22]. The aims of the present study were (i) to compare commonly used primer sets from the literature and a newly designed primer set, and identify the most suitable primer set for metabarcoding of nematodes; (ii) to validate and establish a high-throughput sequencing strategy for nematodes using Illumina paired-end sequencing from individual nematode species as well as bulk DNA from soil. For this, we used single nematodes, mock communities in water and in soil backgrounds, DNA from agricultural fields and from root/rhizosphere samples to validate the primer sets and to test the taxonomic composition of the communities.

## Materials and Methods

### Primer sets

We selected four primer sets for amplicon sequencing of nematodes (S1 Table). The primer set SSU_04F/SSU_22R (SSU) amplifies the V1-V2 region of the 18S rRNA gene (Fig 1) and was recently used to describe assemblages of free-living soil nematodes using the MiSeq platform [17, 23]. We designed a primer set, MMS (MMSF: 5′-GGTGCCAGCAGCCGCGGTA-3′, MMSR: 5′-CTTTAAGT TTCAGCTTTGC-3′) located in the variable region V4-V5 of the 18S rRNA gene (Fig 1). Furthermore, we included the Nemf/18Sr2b primer set (NEM) covering the V6-V8 regions (Fig 1), which has been used to characterize nematode communities from agricultural soils using the Roche 454 platform [18, 21, 24]. Finally, we tested a mitochondrial primer set JB3/JB5 (JB) targeting the I3-M11 region of the COI gene, which has been used to study nematode communities in agricultural field soils and unmanaged flowerbeds in Japan [8].

**Fig 1.**
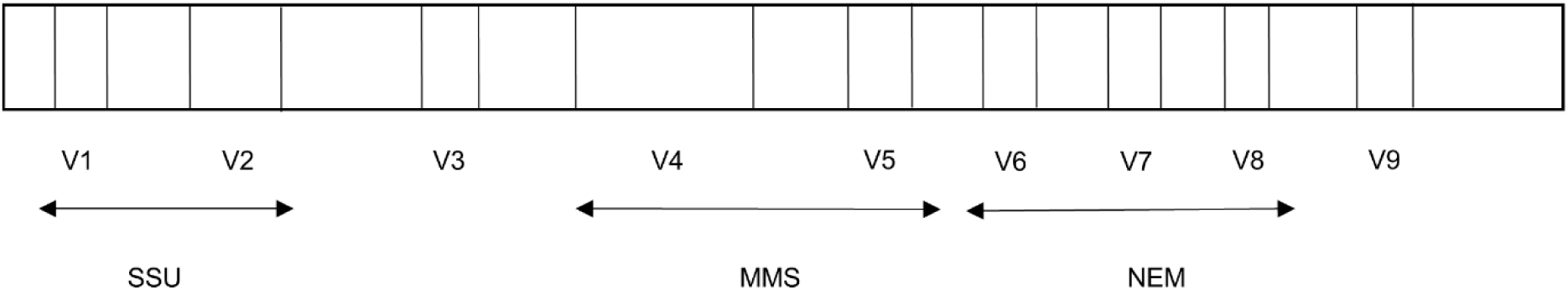
Location of metabarcoding primers targeting variable regions in 18S rRNA gene used in the present study.

### Nematode species, mock communities and root/ rhizosphere soil samples

In order to test the primer sets, we used 22 nematode species obtained from different geographical origins (S2 Table) and artificially assembled 10 mock communities using DNA extracts from these 22 nematodes (named Mock-1, Mock-2 etc.). We combined DNA from the nematode species in different concentrations (S3 Table). We also tested total DNA extracted from soil samples collected from 20 agricultural fields in different parts of Denmark. The field crop history, i.e. the previous crop, stutus, soil type, and pH was recorded (S4 Table). Soil sampling and DNA extraction from the fields were described earlier [24]. Moreover, we spiked 20 nematode mock communities in DNA extracted from soil. We pooled mock community DNA with soil DNA in a 1:1 ratio, referred to as soil-mock communities. Furthermore, we included DNA extracted from washed and freeze-dried root knot nematode (*Meloidogyne incognita*) infected tomato (*Solanum lycopersicum* L.) roots, quinoa (*Chenopodium quinoa* Willd.) roots, maize (*Zea mays* L.) roots/rhizosphere soil, and green bean (*Phaseolus vulgaris* L.) roots/rhizosphere soil.

### DNA extraction, PCR and sequencing strategy

DNA was extracted from 250 mg of the freeze-dried and ground soil samples using the PowerLyzer soil DNA extraction kit (Qiagen, Germany) according to the manufacturer’s instructions, except that samples were homogenized in a Geno/Grinder 2000 at 1500 rpm for 3 x 30 seconds. For the root/rhizosphere soil samples, DNA was extracted from 20 mg of ground material with the DNeasy plant mini kit (Qiagen, Germany).

To amplify target regions, the first PCR was performed in a reaction mixture of 25 μl consisting of 5 µl of Promega 5X reaction buffer, 1.5 µl of MgCl2 (25 mM), 2 µl dNTPs (2.5 mM), 0.5 μl of each primer (10 µM), 0.125 µl of GoTaq Flexi polymerase (5U, Promega Corporation, Madison, USA) and 2 µl of DNA template (approximate 2 ng/µl). PCR cycles for the JB primer combination were 94⁰C for 5 min (94⁰C 1 min, 50⁰C 30 sec, 72⁰C 45 sec) 35 cycles, 72⁰C 10 min, and 4⁰C on hold [25]. Similar PCR cycles were used except that the annealing temperature was 53⁰C for MMS and NEM, and 55⁰C for the SSU primer set [26]. Each of the primer sets of the first PCR (S1 Table) were tagged with the Illumina adaptor overhang nucleotide sequence, for forward primer 5′-TCGTCGGCAGCGTCAGATGTGTATAAGAGACAG-3′ and for reverse primer 5′-GTCTCGTGGGCTCGGAGATGTGTATAAGAGACAG-3′. After this PCR, we pooled and diluted (1:5) the amplicons.

A second PCR was performed for dual indexing. The master mix of this PCR was identical to the first PCR except that 2 μl of DNA template and 2 μl of the different combinations of index primers were used. Each index primer consisted of a sequence specific for Illumina sequencing, a unique 8 bp multiplex identifier and the Illumina adapter overhang sequence. The second PCR was performed with the following cycles: 94⁰C 5 min, (94⁰C 30 sec, 55⁰C 30 sec, 72⁰C 1 min) 13 cycles, 72⁰C 10 min, and 4⁰C on hold. All amplicons were visualized by gel electrophoresis, pooled (approximately equal amounts), precipitated and the pellet dissolved in DNAase free water. Pooled DNA was run on a gel and amplicons were excised and purified using the QIAquick Gel Extraction kit (QIAGEN, Germany) according the manufacturer’s instruction. Finally, the DNA concentrations were measured fluorometrically (Qubit, Thermo Fisher Scientific) and sent for sequencing on an Illumina MiSeq sequencer with PE300 (Eurofins Genomics, Germany).

### Sequence Analysis

The paired end reads obtained from the Illumina MiSeq runs were analyzed using VSEARCH version 2.6 [27]. For joining paired-end reads, we used an overlapping minimum read length of 30 base pairs and reads with quality Phred scores <30 were removed. Internal barcodes, forward and reverse primers, and reads less than 200 base pairs were also excluded. Following this, sequences were dereplicated, screened for chimeras and clustered at 99% similarity level using VSEARCH. Taxonomy assignments for the clustered operational taxonomic units (OTUs) were done using the SILVA 132 reference database [28, 29] for eukaryotes in QIIME using assign_taxonomy.py [30]. Moreover, all nematode OTUs were blasted (≥ 98% similarity) against the NCBI GenBank database to reconfirm their taxonomic assignment. Statistics and data visualization were carried out using the statistical package R.

## Results

### Data characteristics

We successfully obtained sequence reads from 22 individual nematodes species, 20 different mock communities with and without a soil background, 20 different soils and 4 roots/rhizosphere soil samples using the four primer sets. In total, 18.2 million sequence reads were obtained. After quality control, sequence reads were clustered into 320, 17734, 874 and 313 OTUs at 99% similarity for JB, SSU, MMS and NEM primer sets, respectively.

### Sequencing of individual nematode species

For the individual nematode species, we could annotate 10 of the 22 samples to species level and nine to genus level with the JB primer set, whereas three species were not amplified with this primer set (Table 1). Using the SSU primer set, only 15 out of the 22 nematode species were amplified. The MMS primer set amplified all nematodes except *Meloidogyne graminicola* (Table 1). This primer set identified *Meloidogyne* at the genus level, and the other nine nematodes were assigned at species level. The NEM primer set successfully amplified all the nematode species used in our study. *Meloidogyne* species were assigned to genus level and cyst nematodes (*Heterodera carotae* and *H. schachtii*) could only be identified at family level (Table 1). The remaining five nematodes were detected at the species level.

**Table 1.**
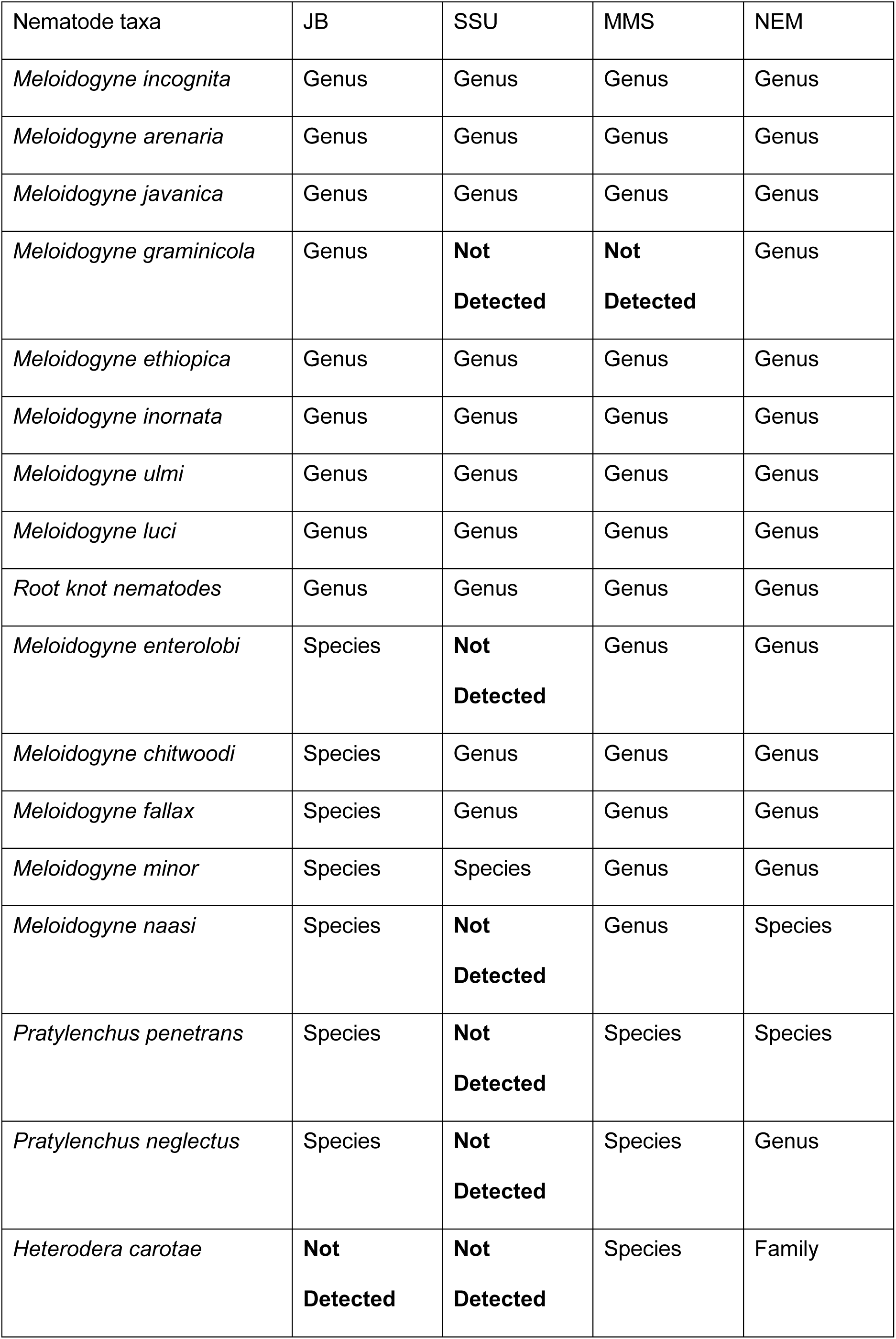
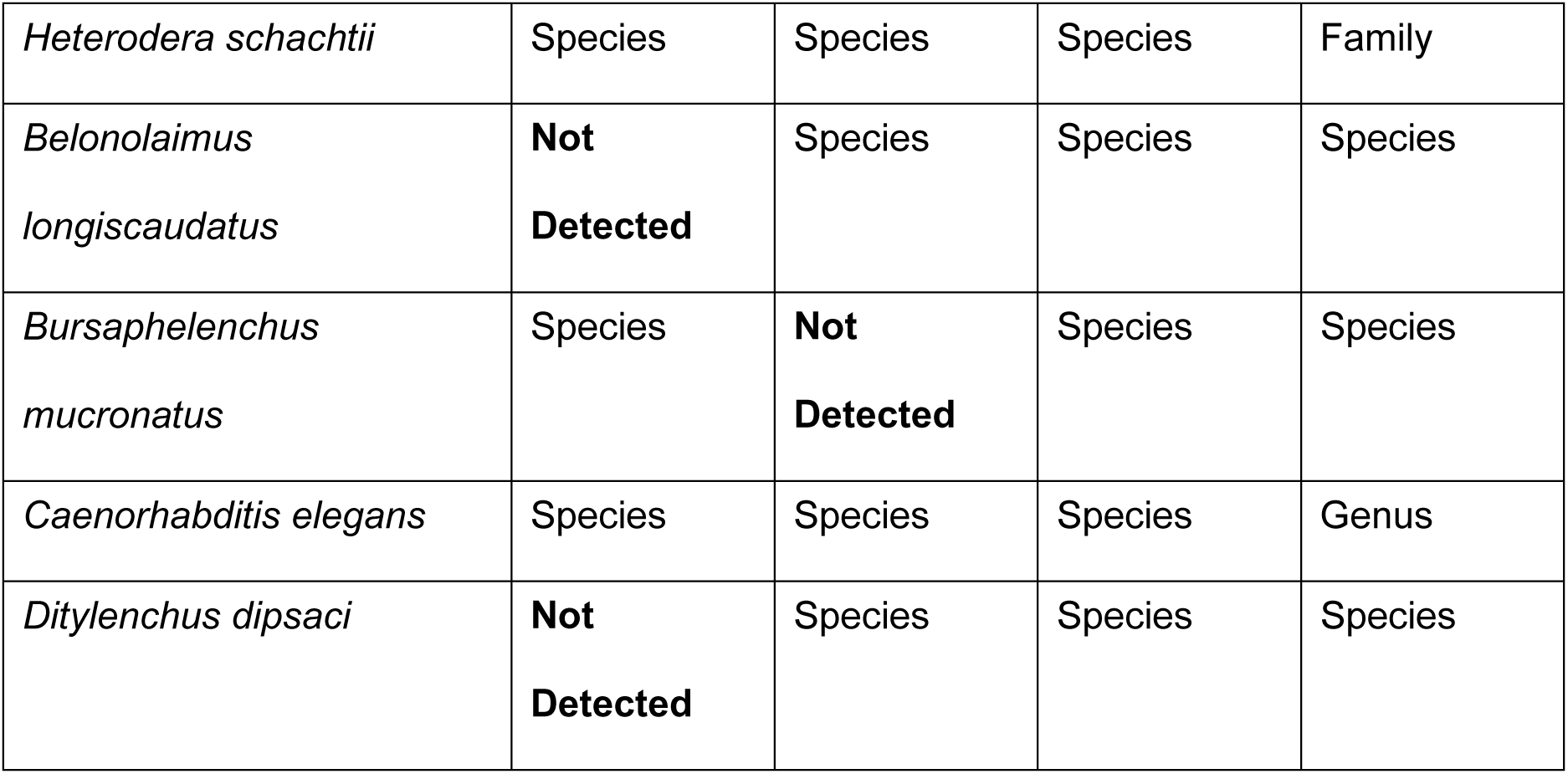
The efficiency of four metabarcoding primer sets in detection of individual nematode species. NCBI Blast tool was used for taxonomic assignments, and top hits with sequence similarities ≥ 99% and coverage 100% were considered for taxonomic assignment at species level.

| Nematode taxa | JB | SSU | MMS | NEM |
| --- | --- | --- | --- | --- |
| *Meloidogyne incognita* | Genus | Genus | Genus | Genus |
| *Meloidogyne arenaria* | Genus | Genus | Genus | Genus |
| *Meloidogyne javanica* | Genus | Genus | Genus | Genus |
| *Meloidogyne graminicola* | Genus | **Not**<br>**Detected** | **Not**<br>**Detected** | Genus |
| *Meloidogyne ethiopica* | Genus | Genus | Genus | Genus |
| *Meloidogyne inornata* | Genus | Genus | Genus | Genus |
| *Meloidogyne ulmi* | Genus | Genus | Genus | Genus |
| *Meloidogyne luci* | Genus | Genus | Genus | Genus |
| *Root knot nematodes* | Genus | Genus | Genus | Genus |
| *Meloidogyne enterolobi* | Species | **Not**<br>**Detected** | Genus | Genus |
| *Meloidogyne chitwoodi* | Species | Genus | Genus | Genus |
| *Meloidogyne fallax* | Species | Genus | Genus | Genus |
| *Meloidogyne minor* | Species | Species | Genus | Genus |
| *Meloidogyne naasi* | Species | **Not**<br>**Detected** | Genus | Species |
| *Pratylenchus penetrans* | Species | **Not**<br>**Detected** | Species | Species |
| *Pratylenchus neglectus* | Species | **Not**<br>**Detected** | Species | Genus |
| *Heterodera carotae* | **Not**<br>**Detected** | **Not**<br>**Detected** | Species | Family |
| *Heterodera schachtii* | Species | Species | Species | Family |
| *Belonolaimus longicaudatus* | **Not Detected** | Species | Species | Species |
| *Bursaphelenchus mucronatus* | Species | **Not Detected** | Species | Species |
| *Caenorhabditis elegans* | Species | Species | Species | Genus |
| *Ditylenchus dipsaci* | **Not Detected** | Species | Species | Species |

### Mock communities and mock communities in soil

In mock communities, JB produced sequences were assigned to genus level within the families Meloidogynidae and Heteroderidae, and to species level within Pratylenchidae and Rhabditidae (Table 2). However, one-third (33%) of the dataset remained unassigned (S1 Fig), and nematodes from Dolichodoridae were not amplified (Table 2). The SSU primer set generated sequences that were assigned to genus level within the Meloidogynidae and Heteroderidae, whereas sequences within Rhabditidae were assigned to species level. The SSU primers failed to amplify nematodes from Pratylenchidae and Dolichodoridae (Table 2). The MMS primer pair generated Meloidogynidae sequences that could be assigned to genus level and for the other three families, Heteroderidae, Dolichodoridae, and Rhabditidae sequences were assigned to species level (Table 2). The NEM primer set was able to amplify and sequence nematodes to the genus level within Meloidogynidae, to species level within Pratylenchidae, Dolichodoridae and Rhabditidae, and to family level within Heteroderidae (Table 2). In the mock communities including diluted DNA of individual nematodes, we observed lower relative abundance of diluted taxa compared to undiluted taxa; however, diluted samples were generally detected in unexpectedly high amounts (S2 and S3 Figs).

**Table 2.** The efficiency of four metabarcoding primers in detection of nematodes in mock communities based on BLAST searches. Only taxonomic assignments appearing in top hits and with sequence similarities ≥ 99% and coverage 100% were considered.

| **Mock communities** | **JB** | **SSU** | **MMS** | **NEM** |
| --- | --- | --- | --- | --- |
| **Meloidogynidae** | Genus | Genus | Genus | Genus |
| **Heteroderidae** | Genus | Species | Species | Family |
| **Pratylenchidae** | Species | **Not detected** | **Not detected** | Species |
| **Dolichodoridae** | **Not detected** | **Not detected** | Species | Species |
| **Rhabditidae** | Species | Species | Species | Species |

In soil-mock combinations, reads from nematodes from the mock communities were generally highly abundant compared to reads from the nematodes derived from the soil background (S5 Table). The JB primer pair only detected nematode families from the mock communities and no additional sequences from the soil background were detected. The SSU primer set was able to detect nematodes belonging to three families (Meloidogynidae, Heteroderidae and Rhabditidae) of the mock communities. Both the MMS and NEM primer sets detected nematodes of the families represented in the mock communities and additionally other nematode families from the spiked soil samples.

### Nematode communities in soil samples

For the JB primer set, 4% and 31% of the total number of sequence reads were classified as Nematoda in soil and plant root/rhizosphere soil samples, respectively, while many sequence reads were unassigned (Fig 2). For the SSU primer set, only 1% of the sequence reads were classified as Nematoda, both in soil and plant root/rhizosphere soil samples (Fig 2). This primer set detected a broad spectrum of other eukaryotes such as fungi, plant, Cercozoa and Charophyta. For the newly designed primer pair (MMSF/MMSR), 17% and 34% of total sequence reads belonged to Nematoda (Fig 2), and for the NEM primer set, 74% and 99% of the total sequences belonged to Nematoda in the soil and plant root/rhizosphere soil samples, respectively (Fig 2).

**Fig 2.**
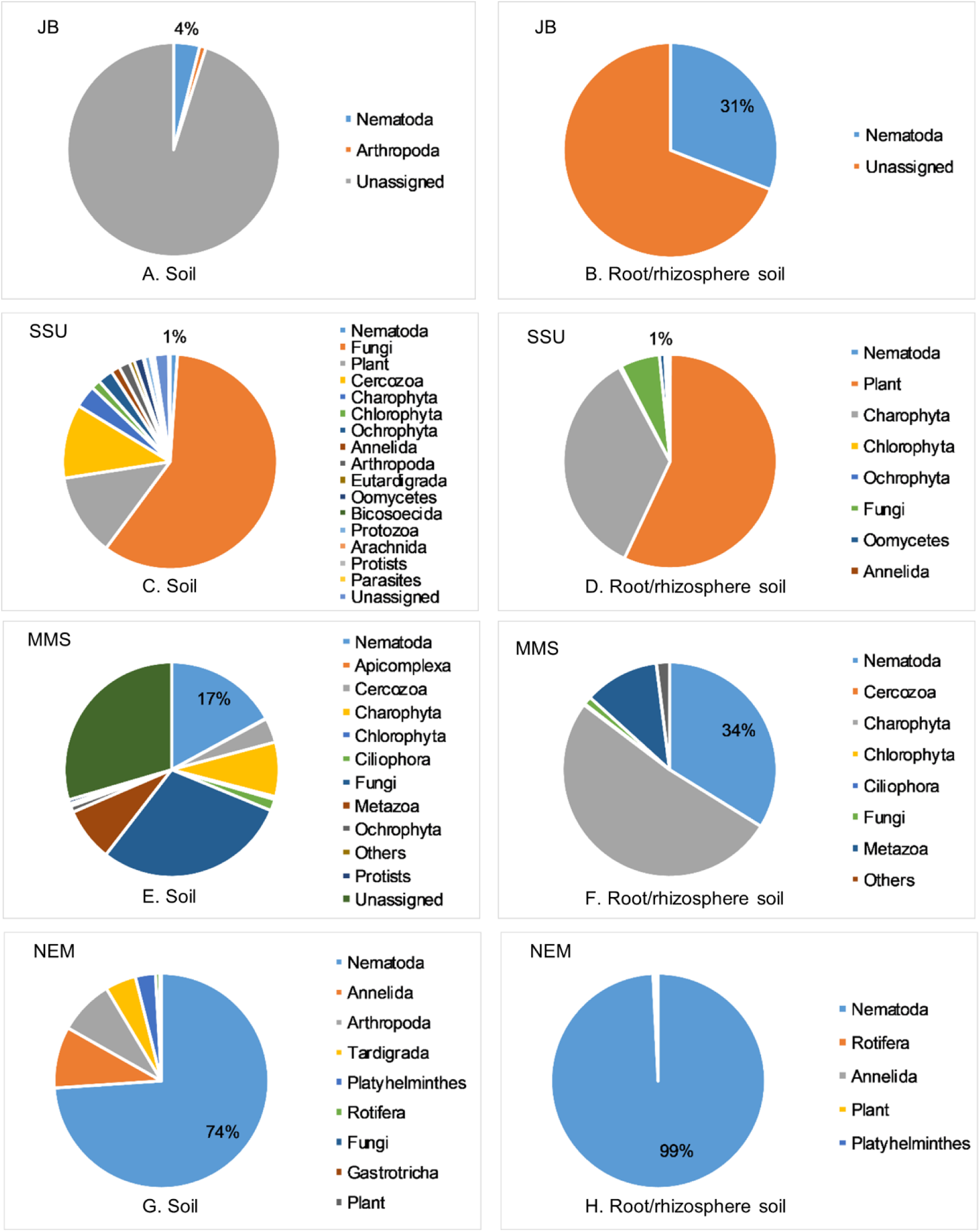
Relative distribution of sequence reads in soil and plant root/rhizosphere soil samples amplified with primer sets JB, SSU, MMS, and NEM; percentage in the blue slide indicates the proportion of sequence reads that were assigned to Nematoda.

We are not presenting any further results for the JB and SSU primer sets due to their poor performance (Fig 2). In the soil samples, NEM and MMS detected a wide range of nematodes from different families (Figs 3 and 4). We recovered 30 nematode families using both the NEM and the MMS primer sets with 6 unique families detected by each primer set (Fig 3). We recorded 14 and 15 different unique genera in soil samples with NEM and MMS primer sets, respectively, and 23 genera were detected by both primer sets (Fig 3). We found 34 and 26 unique nematode species with the NEM and MMS primer set, respectively, while 25 species were detected by both primer sets (Fig 3). We observed that the ability of the two primer sets MMS and NEM to detect nematode families were comparable expect for a limited number of nematode families e.g. Rhabditidae, Trichodoridae, Merliniidae, Heteroderidae (Fig 4 and S6 Table).

**Fig 3.**
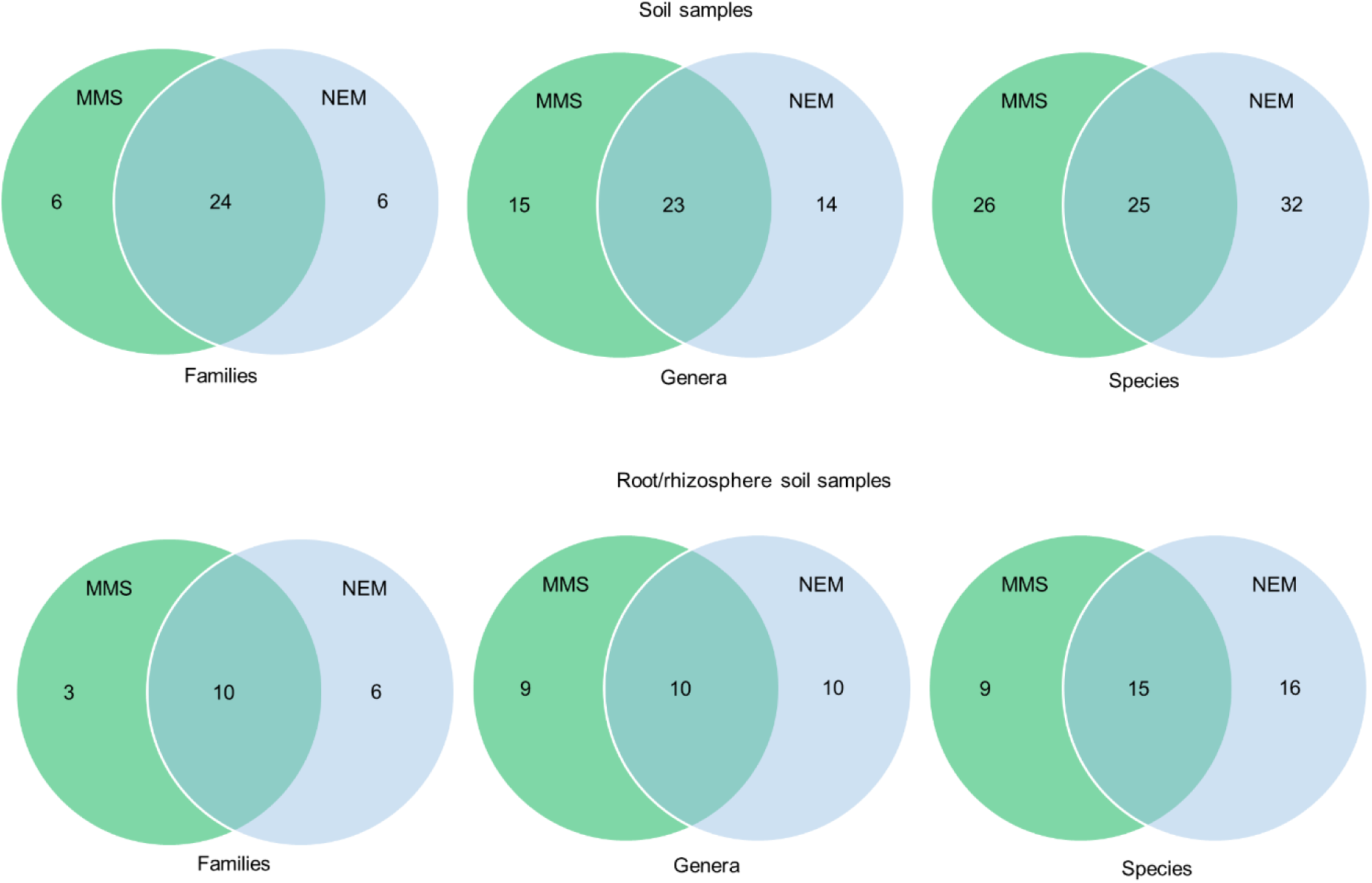
Venn diagrams showing the number of taxa detected in soil samples by the primer sets MMS and NEM. Only taxonomic assignments appearing in top hits of BLAST searches and with sequence similarities ≥ 98% and 100% coverage were considered.

**Fig 4.**
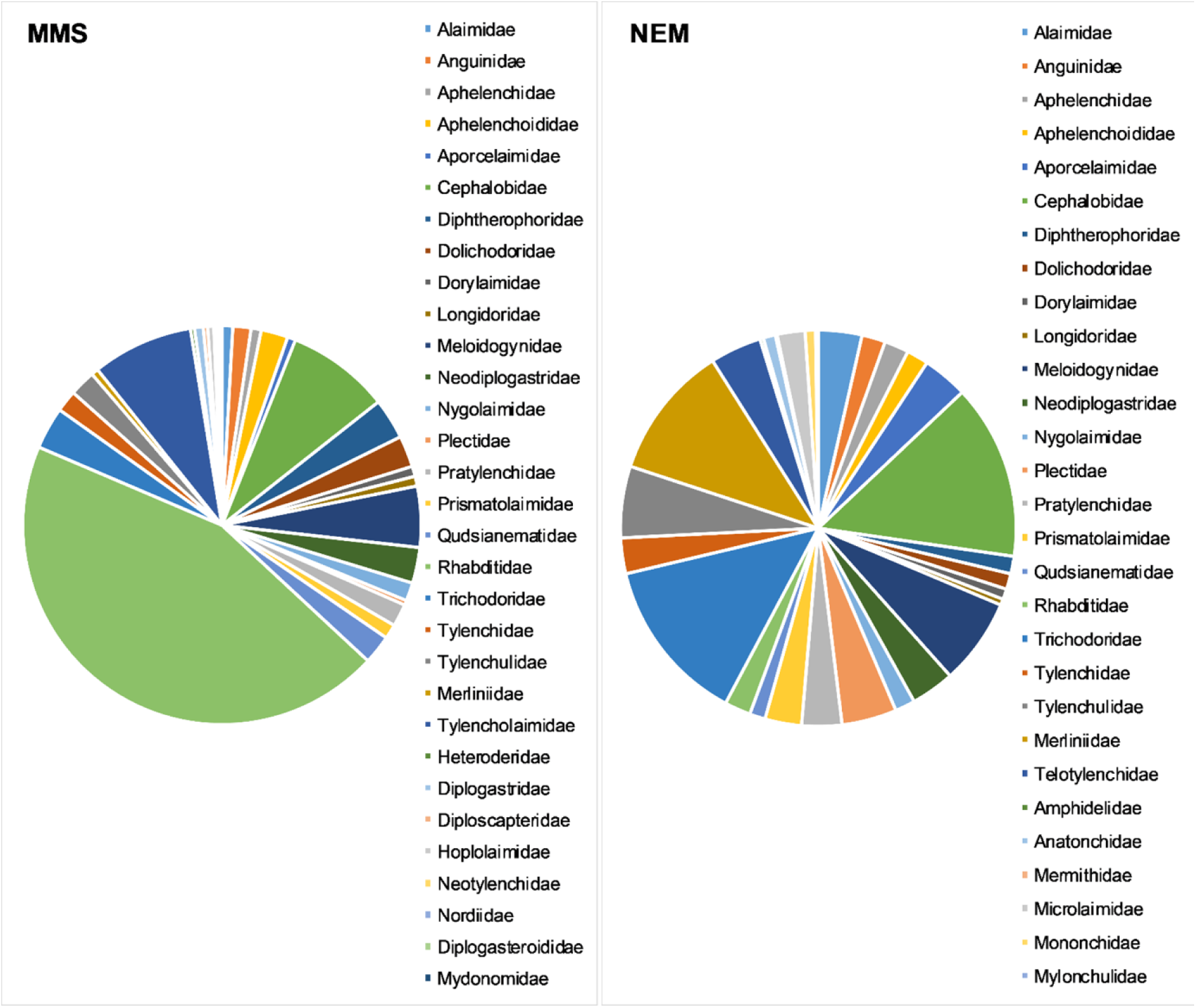
Relative distribution of nematode sequence reads in all soil samples amplified and sequenced with primer sets MMS and NEM. Only taxonomic assignments appearing in top hits of BLAST searches and with sequence similarities ≥ 98% and 100% coverage were considered.

### Nematode communities in plant root/rhizosphere soil samples

In plant root/rhizosphere soil samples, we recovered 16 families by the NEM primers, followed by 13 families using the MMS primer set (Fig 3). Ten families were detected by both primer sets and we recorded 10 and 9 unique genera with NEM and MMS primer sets, respectively, while 10 genera were detected by both primer sets (Fig 3). We detected 16 unique nematode species using the NEM primer set, and 9 unique species were detected using the MMS primers and 15 species were detected by both primer sets. Both primer sets detected a large variation in nematode presence in the samples, and the two primer pairs showed variation in recovered nematode taxa (Fig 5). The quinoa roots and the root knot nematode infected tomato roots were dominated by Meloidogynidae. Both primer sets detected plant parasitic and free-living nematode taxa in green bean and maize root/rhizospheres soils samples.

**Fig 5.**
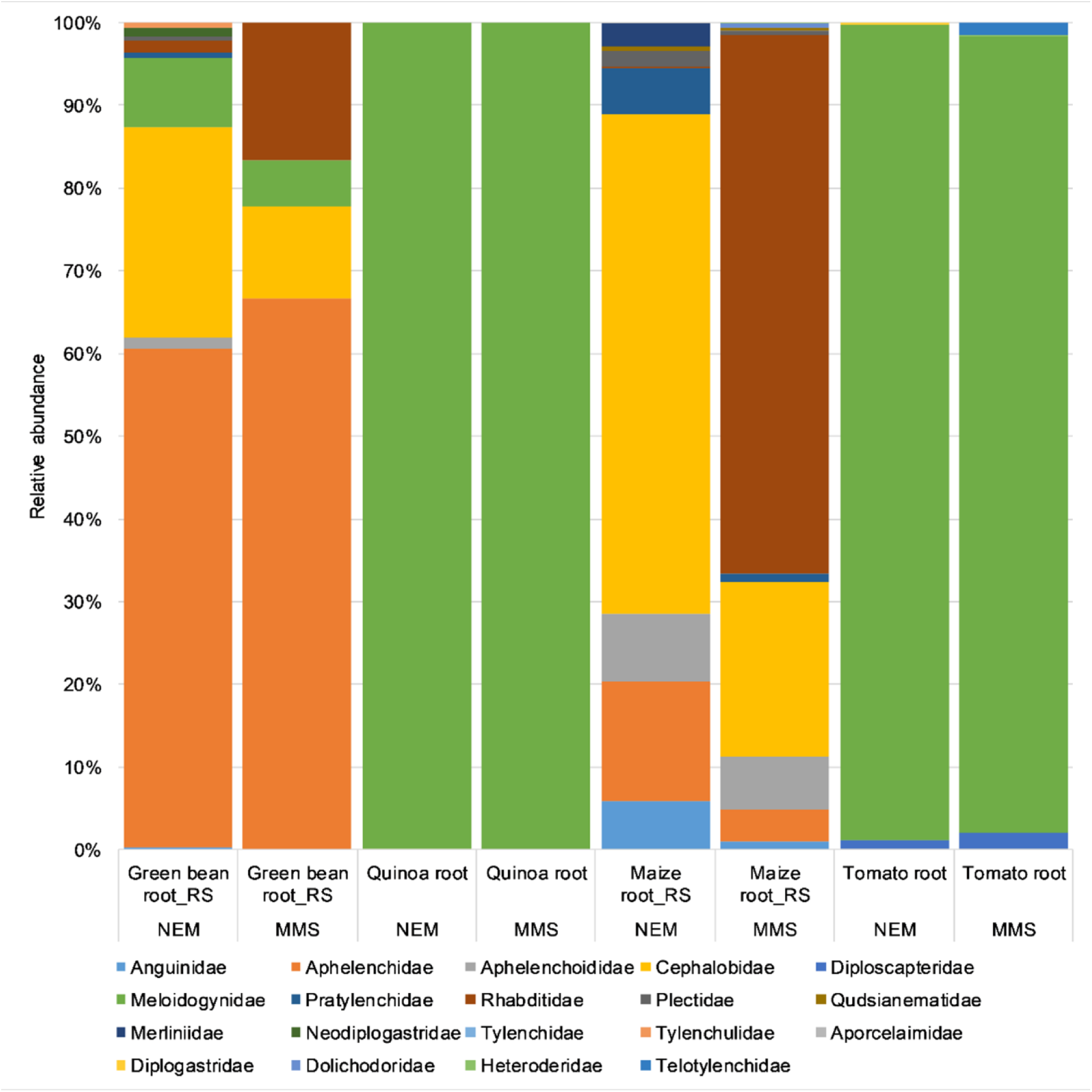
Relative distribution of sequence reads at family rank in plant root/rhizosphere soil samples amplified with primer sets MMS and NEM. Only taxonomic assignments appearing in top hits of BLAST searches and with sequence similarities ≥ 98% and 100% coverage were considered.

## Discussion

Most protocols for nematode metabarcoding include a nematode extraction step to reduce DNA contamination from other soil-living organisms [17–19, 23]. This extraction step may introduce biases as particular nematode taxa or developmental stages are not necessarily extracted at the same efficiency [20]. Furthermore, extraction steps may not be practical when several groups of organisms such as nematodes, fungi and bacteria are studied in the same samples. To overcome these limitations, we previously developed an amplification strategy for 454 pyrosequencing that selectively amplifies nematode DNA from total soil DNA extractions [21]. In the present study, we have adapted amplification strategies for the Illumina MiSeq platform, and we compared different primer sets for their ability to selectively amplify nematode communities.

We observed that the JB primer set only amplified 86% of the individual tested nematode species and did not amplify species that are agronomically important, namely *Heterodera carotae*, *Belonolaimus longicaudatus*, and *Ditylenchus dipsaci*. It has been reported that there are not enough reference sequences of the COI target region in the database for effective species identification [16, 17]. As previously suggested by other researchers, the COI gene has high mutation rates. Hence, the primer sequences are poorly conserved throughout the phylum Nematoda [5, 31]. Based on a study of single nematode species, mock communities, and low number of nematode sequence reads in soil and root/rhizosphere soil samples, we found that the JB primer set targeting the I3-M11 partition of the COI gene is not suitable for nematode metabarcoding.

In a recent study, the SSU ribosomal DNA marker (SSU_04F/SSU_22R) outperformed the mitochondrial marker (JB3/JB5GED) in terms of nematode species and genus level detection [17, 23]. However, in our study, important nematode species were not amplified and detected by the SSU primer set. Moreover, the amplification strategy using the SSU primer set only resulted in 1% Nematoda reads from soil and plant root/rhizosphere soil DNA samples. Our results corroborates a recent study in which the NF1/18Sr2b primer set provided better taxonomic resolutions compared to the SSU_04F/SSU_22R marker [17]. In other studies, this SSU primer set was found to amplify a large portion of non-nematode reads of environmental marine sediment samples [32–35]. Therefore, this primer set was not considered suitable for nematode diversity studies of environmental samples without an initial nematode extraction step.

Results from the analysis of the individual nematodes showed that better taxonomic resolution was achieved with MMS, which targets the V4-V5 region of 18S rRNA gene, compared to JB and SSU primer sets. The efficiency of this primer set was further confirmed using the mock communities as it was able to detect all the nematode taxa in the mock communities, also in a soil background. MMS detected a high diversity of the nematode communities in soil samples, suggesting that this newly designed primer set is well suited for studies of plant parasitic and free-living nematodes. This primer set was also able to detect many nematode families in the plant root/rhizosphere soil samples. Based on these observations, this newly designed MMS primer set is efficient for studies of soil nematode communities, and it clearly outperforms the JB and SSU primer sets.

The NEM primer set was previously developed for the 454-sequencing platform using a semi-nested PCR approach. However, in the present study, the second PCR in the nested PCR was omitted, which resulted in a larger PCR product (500bp) including the V6, V7, and V8 regions of the 18S ribosomal RNA gene. All individual nematode species in our study were identified using NEM, and all nematode families in the mock communities were detected. In addition, NEM detected a range of diverse nematode taxa in the different soils, reflecting the different crop species that had been grown in the soils, and the different soil parameters. The NEM primer set amplified more nematode taxa in the root/rhizosphere soil samples compared to all the other primer sets tested. This primer set amplified almost 100% nematode DNA in the presence of plant DNA, which indicates that this primer set is highly nematode specific.

Although we detected fewer sequence reads using both MMS and NEM primer sets when we used diluted templates in the mock communities, the read counts were not reduced quantitatively. The reason for this is not known.

Sequence reads from taxa that belong to the family Rhabditidae were much more prevalent in the MMS than in the NEM-generated data set. This discrepancy is probably due to a three-nucleotide mismatch between 18Sr2b primer of the NEM primer set and the Rhabditidae DNA template. It was reported that the reverse primer sequence (18Sr2b) failed to amplify several Rhabditidae species [36]. In a recent study, a modified version of the primer set Nemf/18Sr2b, named as NemFopt/18Sr2bRopt, was constructed by adding extra nucleotides and by including degenerate bases in both the forward and reverse primer to improve GC content and shift the reverse primer into a more conserved region of Nematoda [18]. The MMS primer set could overcome problems associated with detecting Rhabditidae species. Multiple sequence alignments (S4 Fig), detection of a higher number of taxa belonging to Rhabditidae and the greater relative abundance of Rhabditidae in soil samples in our sequence data confirmed that the MMS primer set efficiently detects Rhabditiae.

The MMS primer set did not detect nematode taxa of the families Aporcelaimidae, Diplogastridae, Merliniidae, Neodiplogastridae, Tylenchidae, and Tylenchulidae in root/rhizosphere soil samples, although this group of nematode taxa was detected in our soil samples. This fact could be due to the competition in primer annealing between nematode and plant DNA templates in root/rhizosphere soil samples. The NEM primer set was not able to detect the families Heteroderidae, Dolichodoridae, Telotylenchidae in the root/rhizosphere soil samples. We observed that the NEM primer set could not detect *Heterodera carotae, H. schachtii* and *Globodera* spp. neither at genus nor at species level in individual nematode species, mock communities, and mock communities in soil. On the contrary, the MMS primer set was found to be efficient in detecting nematodes belonging to Heteroderidae.

We propose to use both primer sets (MMS and NEM) for identification of nematode communities on DNA extracted directly from soil. Together, these two primer sets cover more than 1000 bp of the 18S rRNA gene and they capture a substantial range of the variable regions (V4, V5, V6, V7, and V8) of the 18S rRNA gene in nematodes. Moreover, the assignment of lower Linnaean taxonomies (genus, species) to sequence reads is a very crucial step in the use of DNA markers for biodiversity assessment. We conclude that our two primer sets (MMS and NEM) complement each other in detecting nematode families and can efficiently detect nematodes at the genus level and in some cases at species level.

## Acknowledgement

We are grateful to Andrea M. Skantar, United State Department of Agriculture; Barbara Geric Stare, Sasa Sirca and Gregor Urek, Department of Plant Protection, Agricultural Institute of Slovenia for contributing DNA samples of nematode species. We would like to thank Susana Santos for data visualization. We would also like to thank Mathilde Schiøtt Dige and Simone Ena Rasmussen for their laboratory technical assistance.

## Supporting information

**S1 Table.**
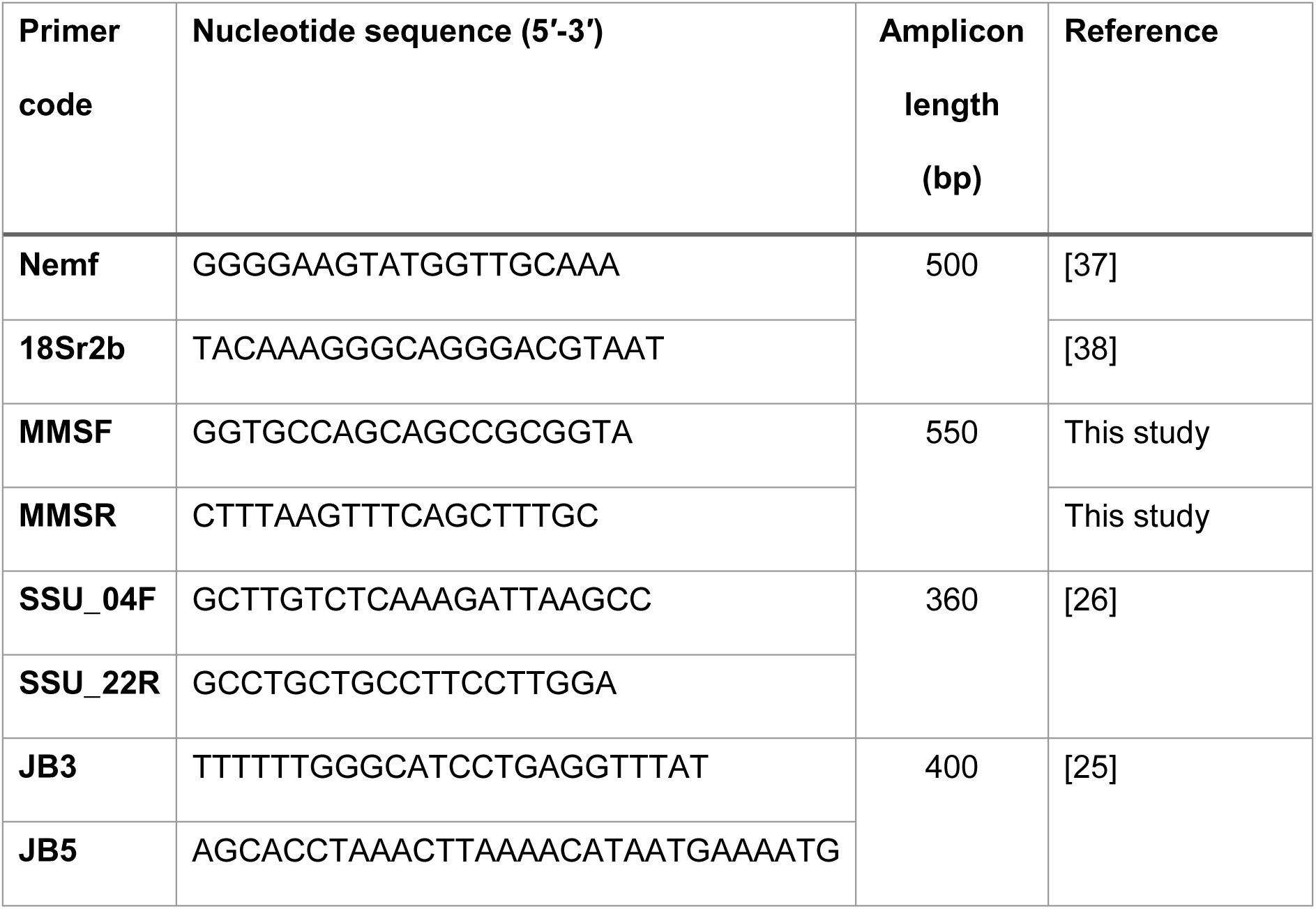
Metabarcoding primer sets used in the present study.

**S2 Table.** List of individual nematode species, DNA extraction method and order of the species used in the study.

|  | Nematode taxa | DNA extraction method | Order |
| --- | --- | --- | --- |
| 1. | *Meloidogyne incognita* | Qiasafe | Tylenchida |
| 2. | *Meloidogyne arenaria* | Qiasafe | Tylenchida |
| 3. | *Meloidogyne nassi* | DNeasy | Tylenchida |
| 4. | *Meloidogyne minor* | Qiasafe | Tylenchida |
| 5. | *Meloidogyne javanica* | Worm lysis buffer | Tylenchida |
| 6. | *Meloidogyne graminicola* | Worm lysis buffer | Tylenchida |
| 7. | *Meloidogyne fallax* | Qiasafe | Tylenchida |
| 8. | *Meloidogyne chitwoodi* | DNeasy | Tylenchida |
| 9. | *Meloidogyne ulmi* | DNeasy | Tylenchida |
| 10. | Mixed *Meloidogyne* spp. | Worm lysis buffer | Tylenchida |
| 11. | *Meloidogyne enterolobi* | Worm lysis buffer | Tylenchida |
| 12. | *Meloidogyne inornata* | Worm lysis buffer | Tylenchida |
| 13. | *Meloidogyne ethiopica* | Worm lysis buffer | Tylenchida |
| 14. | *Meloidogyne luci* | Worm lysis buffer | Tylenchida |
| 15. | *Belonolaimus longicaudatus* | Qiasafe | Tylenchida |
| 16. | *Pratylenchus penetrans* | DNeasy | Tylenchida |
| 17. | *Pratylenchus neglectus* | Worm lysis buffer | Tylenchida |
| 18. | *Heterodera schachtii* | Qiasafe | Tylenchida |
| 19. | *Heterodera carotae* | Qiasafe | Tylenchida |
| 20. | *Caenorhabditis elegans* | DNeasy | Rhabditida |
| 21. | *Ditylenchus dipsaci* | Worm lysis buffer | Tylenchida |
| 22. | *Bursaphelenchus mucronatus* | Worm lysis buffer | Aphelenchida |

**S3 Table.** Composition of mock communities used in the study.

|  | Nematode taxa | Nematode taxa | Nematode taxa | Nematode taxa | Nematode taxa | Nematode taxa |
| --- | --- | --- | --- | --- | --- | --- |
| **Mock1** | *Meloidogyne hapla* | *Globodera* spp. | *Belonolaimus longicaudatus* | *Pratylenchus penetrans* | *Heterodera carotae* | *Caenorhabditis elegans* |
| **Mock2** | *Meloidogyne incognita* | *Globodera* spp. | *B. longicaudatus* | *P. penetrans* | *Heterodera schachtii* | *C. elegans* |
| **Mock3** | *Meloidogyne arenaria* | *Globodera* spp. | *B. longicaudatus* | *P. penetrans* | *H. carotae* | *C. elegans* |
| **Mock4** | *Meloidogyne minor* | *Globodera* spp | *B. longicaudatus* | *P. penetrans* | *H.schachtii* | *C. elegans* |
| **Mock5** | *Meloidogyne fallax* | *Globodera* spp | *B. longicaudatus* | *P. penetrans* | *H. carotae* | *C. elegans* |
| **Mock6** | *Meloidogyne chitwoodi* | *Globodera* spp | *B. longicaudatus* | *P. penetrans* | *H.schachtii* | *C. elegans* |
| **Mock7** | *Meloidogyne ulmi*+<br>*Meloidogyne nassi* | *Globodera* spp | *B. longicaudatus* | *P. penetrans* | *H. carotae* | *C. elegans* |
| **Mock8** | *Meloidogyne inornata*<br>+Mixed *Meloidogyne* spp. | *Globodera* spp | *B. longicaudatus* | *P. penetrans* | *H.schachtii* | *C. elegans* |
| **Mock9** | *Meloidogyne ethiopica*<br>+Mixed *Meloidogyne* spp. | *Globodera* spp | *B. longicaudatus* | *P. penetrans* | *H. carotae* | *C. elegans* |
| **Mock10** | *Meloidogyne luci*+<br>*Meloidogyne inornata* | *Globodera* spp | *B. longicaudatus* | *P. penetrans* | *H.schachtii* | *C. elegans* |
| **Mock11** | *Meloidogyne hapla* | *Globodera* spp | *B. longicaudatus* | *P. penetrans* | *H. carotae* | *C. elegans* |
| **Mock12** | *Meloidogyne incognita* | *Globodera* spp | *B. longicaudatus* | *P. penetrans* | *H.schachtii* | *C. elegans* |
| **Mock13** | *Meloidogyne arenaria* | *Globodera* spp | *B. longicaudatus* | *P. penetrans* | *H. carotae* | *C. elegans* |
| **Mock14** | *Meloidogyne minor* | *Globodera* spp | *B. longicaudatus* | *P. penetrans* | *H.schachtii* | *C. elegans* |
| **Mock15** | *Meloidogyne fallax* | *Globodera* spp | *B. longicaudatus* | *P. penetrans* | *H. carotae* | *C. elegans* |
| **Mock16** | *Meloidogyne chitwoodi* | *Globodera* spp | *B. longicaudatus* | *P. penetrans* | *H. schachtii* | *C. elegans* |
| **Mock17** | Mixed *Meloidogyne* spp. + *Meloidogyne nassi*+ *Meloidogyne inornata* | *Globodera* spp | *B. longicaudatus* | *P. penetrans* | *H. carotae* | *C. elegans* |
| **Mock18** | *Meloidogyne inornata* | *Globodera* spp | *B. longicaudatus* | *P. penetrans* | *H. schachtii* | *C. elegans* |
| **Mock19** | *Meloidogyne ethiopica* | *Globodera* spp | *B. longicaudatus* | *P. penetrans* | *H. carotae* | *C. elegans* |
| **Mock20** | *Meloidogyne luci* | *Globodera* spp | *B. longicaudatus* | *P. penetrans* | *H. schachtii* | *C. elegans* |

Blue font indicates 1:10 dilution of template DNA

**S4 Table.** Cropping history and soil properties of twenty different soils used in the study.

| Soil ID | Previous crops | Year of sampling | Status | Soil type | pH |
| --- | --- | --- | --- | --- | --- |
| **Soil-1** | Carrot | 2012 | NA | NA | NA |
| **Soil-2** | Strawberry | 2012 | NA | NA | NA |
| **Soil-3** | Spinach | 2014 | NA | NA | NA |
| **Soil-4** | Maize | 2018 | Conventional | *clayey sand* | 6.0 |
| **Soil-5** | Beans | 2018 | Conventional | *clayey sand* | 5.9 |
| **Soil-6** | Triticale | 2013 | Conventional | *heavy clay* | 5.8 |
| **Soil-7** | Corn | 2014 | Conventional | *heavy clay* | 6.3 |
| **Soil-8** | Iceberg lettuce | 2018 | Conventional | *clayey sand* | 6.3 |
| **Soil-9** | Rye | 2013 | Conventional | *heavy clay* | 5.9 |
| **Soil-10** | Rye | 2014 | Organic | *clay* | 6.0 |
| **Soil-11** | Barley | 2013 | Organic | *clayey sand* | 5.2 |
| **Soil-12** | Barley | 2014 | Conventional | *heavy clay* | 6.5 |
| **Soil-13** | Wheat | 2013 | Conventional | *heavy clay* | 6.5 |
| **Soil-14** | Wheat | 2014 | Conventional | *coarse sand* | 6.1 |
| **Soil-15** | Potato | 2013 | Conventional | *heavy clay* | 5.4 |
| **Soil-16** | Potato | 2014 | Conventional | *heavy clay* | 5.5 |
| **Soil-17** | Clover | 2013 | Organic | *heavy clay* | 5.7 |
| **Soil-18** | Clover | 2014 | Organic | *coarse sand* | 6.1 |
| **Soil-19** | Oat | 2013 | Organic | *heavy clay* | 5.9 |
| **Soil-20** | Oat | 2014 | Organic | *heavy clay* | 6.0 |

NA indicates not analysed

**S5 Table.** The efficiency of four metabarcoding primers in the detection of nematodes from different families in twenty different soil mock communities.

| **Families** | **JB** | **SSU** | **MMS** | **NEM** |
| --- | --- | --- | --- | --- |
| **Alaimidae** | ND | D | D | D |
| **Amphidelidae** | ND | ND | ND | D |
| **Anatonchidae** | ND | ND | ND | D |
| **Anguinidae** | ND | ND | D | D |
| **Aphelenchidae** | ND | ND | D | D |
| **Aphelenchoididae** | ND | ND | D | D |
| **Aporcelaimidae** | ND | ND | D | D |
| **Axonolaimidae** | ND | D | ND | ND |
| **Bastianiidae** | ND | D | ND | ND |
| **Cephalobidae** | ND | D | D | D |
| **Diplopeltidae** | ND | D | ND | ND |
| **Diphterophoridae** | ND | D | D | D |
| **Diplogasteridae** | ND | ND | ND | D |
| **\*Dolichodoridae** | ND | ND | D | D |
| **Dorylaimidae** | ND | ND | D | D |
| **\*Heteroderidae** | D | D | D | D |
| **Longidoridae** | ND | ND | ND | D |
| **\*Meloidogynidae** | D | D | D | D |
| **Merliniidae** | ND | ND | ND | D |
| **Mermithidae** | ND | ND | ND | D |
| **Microlaimidae** | ND | ND | ND | D |
| **Monhysteridae** | ND | D | ND | ND |
| **Mononchidae** | ND | ND | ND | D |
| **Mydonomidae** | ND | ND | D | ND |
| **Mylonchulidae** | ND | ND | ND | D |
| **Neodiplogastridae** | ND | ND | D | D |
| **Nygolaimidae** | ND | ND | D | D |
| **Plectidae** | ND | D | ND | D |
| **\*Pratylenchidae** | D | ND | D | D |
| **Prismatolaimidae** | ND | D | D | D |
| **Qudsianematidae** | ND | ND | D | D |
| **\*Rhabditidae** | D | D | D | D |
| **Telotylenchidae** | ND | ND | D | D |
| **Trichodoridae** | ND | D | ND | D |
| **Tylenchidae** | ND | ND | D | D |
| **Tylenchulidae** | ND | ND | D | D |
| **Unassigned** | Above 70% | - | Negligible | Negligible |

Here, D denotes amplified and detected, ND denotes not detected, \* denotes the families were abundant in soil-mock samples.

**S6 Table.** Efficiency of four metabarcoding primers in detection of nematodes from twenty different soils at lower taxonomic rank than family level.

| Taxa | JB | SSU | MMS | NEM |
| --- | --- | --- | --- | --- |
| **Achromadoridae** | ND | D | ND | ND |
| *Achromadora* sp. |  | G |  |  |
| *Achromadora ruricola* |  | S |  |  |
| **Alaimidae** | ND | D | D | D |
| *Alaimus* sp. |  | G | G | G |
| *Alaimus arcuatus* |  |  |  | S |
| *Alaimus parvus* |  |  |  | S |
| **Anatonchidae** | ND | ND | ND | D |
| *Anatonchus tridentatus* |  |  |  | S |
| **Anguinidae** | ND | ND | D | D |
| *Ditylenchus* sp. |  |  | G | G |
| *Ditylenchus dipsaci* |  |  |  | S |
| *Ditylenchus destructor* |  |  |  | S |
| **Aphelenchidae** | ND | ND | D | D |
| *Aphelenchus avenae* |  |  | S | S |
| **Aphelenchoididae** | ND | ND | D | D |
| *Aphelenchoides* sp. |  |  | G |  |
| *Aphelenchoides parietinus* |  |  | S | S |
| *Aphelenchoides bicaudatus* |  |  | S | S |
| *Seinura demani* |  |  |  | S |
| **Aporcelaimidae** | ND | ND | D | D |
| *Aporcelaimellus* sp. |  |  | G | G |
| Bastianiidae | ND | D | ND | ND |
| *Bastiania gracilis* |  | S |  |  |
| Cephalobidae | ND | D | D | D |
| *Acrobeles complexus* |  | S | S | S |
| *Acrobeles ctenocephalus* |  |  |  | S |
| *Acrobeles ciliatus* |  | S |  | S |
| *Acrobeloides* sp. |  | G | G | G |
| *Acrobeloides thornei* |  | S |  |  |
| *Acrobeloides apiculatus* |  | S |  |  |
| *Acrobeloides varius* |  |  |  | S |
| *Chiloplacus propinquus* |  | S | S |  |
| *Eucephalobus* sp. |  |  |  | G |
| *Eucephalobus striatus* |  | S | S | S |
| *Eucephalobus oxyuroides* |  | S | S |  |
| *Heterocephalobus elongatus* |  |  | S |  |
| *Pseudacrobeles* sp. |  |  |  | G |
| Diplopeltidae | ND | D | ND | ND |
| *Cylindrolaimus communis* |  | S |  |  |
| Diphtherophoridae | ND | D | D | D |
| *Diphtherophora* sp. |  | G | G | G |
| *Diphtherophora communis* |  | S | S |  |
| *Tylolaimophorus typicus* |  | S |  |  |
| **Diplogasteridae** | ND | D | ND | ND |
| *Butlerius butleri* |  | S |  |  |
| **Diplogasteroididae** | ND | ND | D | ND |
| *Diplogasteroides sp.* |  |  | G |  |
| **Diploscapteridae** | ND | ND | D | ND |
| *Diploscapter sp.* |  |  | G |  |
| **Dolichodoridae** | ND | ND | D | D |
| *Merlinius sp* |  |  |  | G |
| *Merlinius nanus* |  |  | S |  |
| **Dorylaimidae** | ND | ND | D | D |
| *Mesodorylaimus bastiani* |  |  | S |  |
| *Prodorylaimus sp.* |  |  |  | G |
| *Thonus circulifer* |  |  | S | S |
| **Heteroderidae** | D | ND | D | ND |
| *Globodera pallida* | S |  | S |  |
| *Globodera rostochiensis* | S |  |  |  |
| **Hoplolaimidae** | ND | ND | D | ND |
| *Helicotylenchus minzi* |  |  | S |  |
| **Longidoridae** | ND | ND | D | D |
| *Longidorus sp.* |  |  | G |  |
| *Longidorus attenuatus* |  |  |  | S |
| **Meloidogynidae** | D | D | D | D |
| *Meloidogyne sp.* | G | G | G | G |
| *Meloidogyne hapla* |  |  | S | S |
| Merliniidae | ND | ND | *D | ND |
| Microlaimidae | ND | ND | ND | D |
| *Prodesmodora circulata* |  |  |  | S |
| Mylonchulidae | ND | ND | ND | D |
| *Mylonchulus hawaiiensis* |  |  |  | S |
| Mononchidae | ND | ND | ND | D |
| *Clarkus papillatus* |  |  |  | S |
| Monhysteridae | ND | D | ND | ND |
| *Eumonhystera* sp. |  | G |  |  |
| *Eumonhystera vulgaris* |  | S |  |  |
| *Eumonhystera hungarica* |  | S |  |  |
| *Geomonhystera* sp. |  | G |  |  |
| Mydonomidae | ND | ND | D | ND |
| *Dorylaimoides micoletzkyi* |  |  | S |  |
| Neodiplogastridae | ND | ND | D | D |
| *Mononchoides americanus* |  |  | S |  |
| *Pristionchus* sp. |  |  | G | G |
| *Pristionchus lheritieri* |  |  |  | S |
| Neotylenchidae | ND | ND | D | ND |
| *Rubzovinema* sp. |  |  | G |  |
| Nordiidae | ND | ND | D | ND |
| *Pungentus* sp. |  |  | G |  |
| **Nygolaimidae** | ND | ND | D | D |
| *Nygolaimus brachyuris* |  |  | S | S |
| **Plectidae** | ND | D | ND | D |
| *Plectus* sp. |  |  |  | G |
| *Plectus minimus* |  | S |  |  |
| *Plectus aquatilis* |  | S |  |  |
| *Anaplectus porosus* |  | S |  | S |
| *Tylocephalus auriculatus* |  | S |  |  |
| *Wilsonema otophorum* |  | S |  |  |
| **Pratylenchidae** | D | ND | D | D |
| *Pratylenchus thornei* |  |  | S | S |
| *Pratylenchus crenatus* |  |  | S | S |
| *Pratylenchus penetrans* |  |  | S | S |
| *Pratylenchus neglectus* |  |  | S | S |
| *Pratylenchus fallax* | S |  |  |  |
| **Prismatolaimidae** | ND | D | D | D |
| *Prismatolaimus* sp. |  |  |  | G |
| *Prismatolaimus dolichurus* |  | S | S |  |
| *Prismatolaimus intermedius* |  | S | S |  |
| **Qudsianematidae** | ND | ND | D | D |
| *Microdorylaimus miser* |  |  |  | S |
| *Ecumenicus monohystera* |  |  | S | S |
| **Rhabditidae** | D | ND | D | D |
| *Rhabditis* sp. |  |  | G | G |
| *Pelodera* sp. |  |  | G |  |
| *Pelodera teres* |  |  |  | S |
| *Pellioiditis* sp. |  |  |  | G |
| *Pellioiditis marina* | S |  |  |  |
| *Mesorhabditis* sp. |  |  | G |  |
| *Mesorhabditis belari* |  |  | S |  |
| *Caenorhabditis elegans* | S |  |  |  |
| Steinernematidae | ND | D | ND | ND |
| *Steinernema affine* |  | S |  |  |
| Telotylenchidae | ND | D | D | D |
| *Tylenchorhynchus maximus* |  |  | S | S |
| *Tylenchorhynchus dubius* |  |  | S | S |
| *Tylenchorhynchus teeni* |  | S |  |  |
| Trichodoridae | ND | D | D | D |
| *Trichodorus primitivus* |  | S | S | S |
| *Trichodorus viruliferus* |  | S |  |  |
| *Paratrichodorus pachydermus* |  |  | S | S |
| *Paratrichodorus allius* |  | S |  |  |
| Tylenchidae | ND | ND | D | D |
| *Filenchus* sp. |  |  |  | G |
| *Filenchus aquilonius* |  |  |  | S |
| *Coslenchus turkeyensis* |  |  | S |  |
| ***Basiria* sp.** |  |  |  | G |
| ***Basiria duplexa*** |  |  | S |  |
| **Tylenchulidae** | ND | ND | D | D |
| ***Paratylenchus* sp.** |  |  |  | G |
| ***Paratylenchus conicephalus*** |  |  |  | S |
| ***Paratylenchus similis*** |  |  |  | S |
| ***Paratylenchus nanus*** |  |  | S |  |
| ***Paratylenchus projectus*** |  |  | S |  |
| **Tylencholaimidae** | ND | ND | D | ND |
| ***Tylencholaimus* sp.** |  |  | G |  |

Here, G: genus level detection, S: species level detection, D: detected at lower taxonomic level, \*D: detected at family level only, ND: not amplified and detected at any taxonomic level. Classification was based on Nemaplex database (http://nemaplex.ucdavis.edu/ Accessed on 11/11/2019); NCBI Blast tool was used for taxonomic assignments, and top hits with sequence similarities $\geq 99\%$ and coverage 100% were considered for taxonomic assignment at species level.

**S1 Fig.**
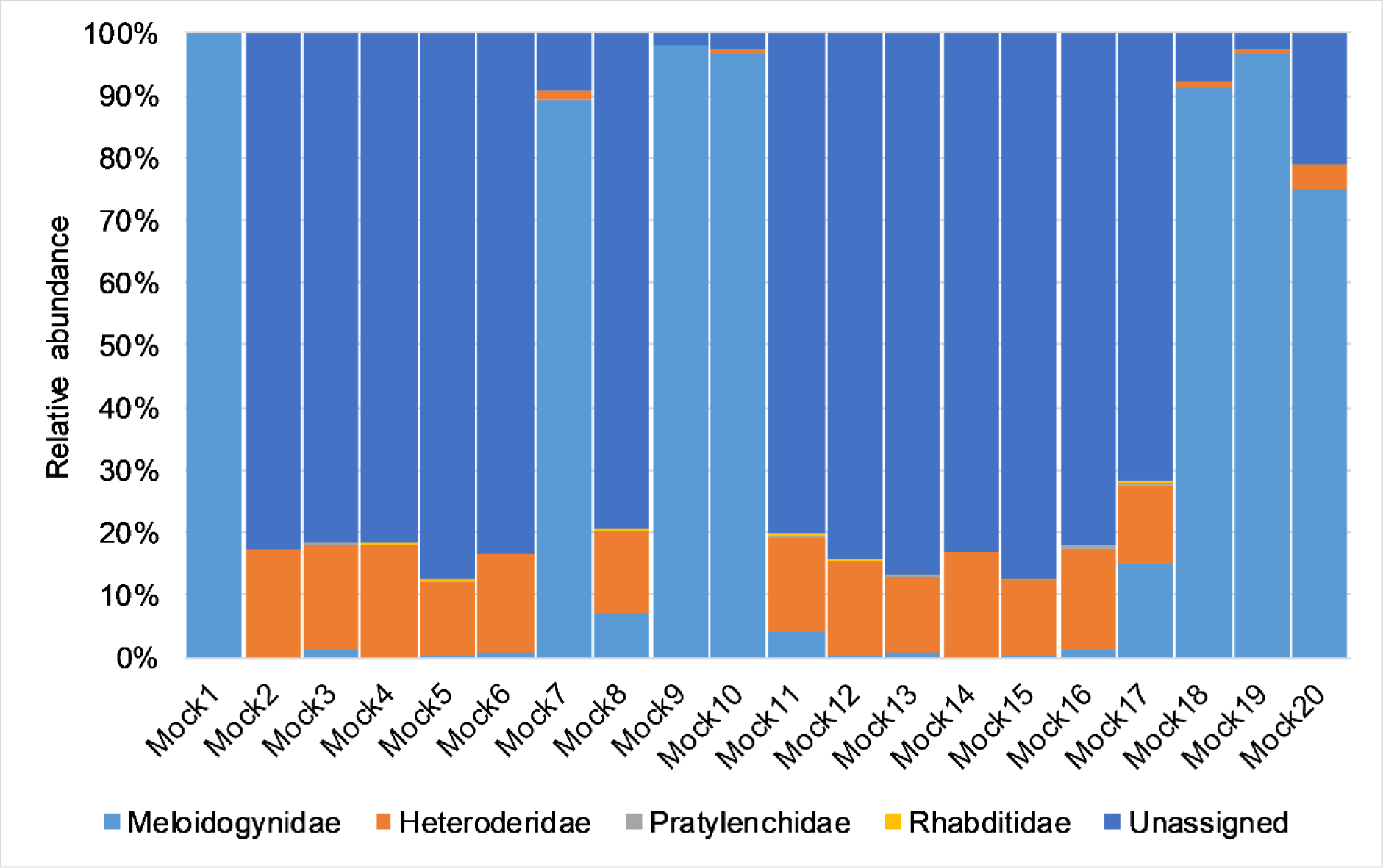
Relative abundance of sequence reads at family rank in mock samples amplified and sequenced using JB primer set. Only taxonomic assignments appearing in top hits and with sequence similarities ≥ 99% and coverage 100% were considered.

**S2 Fig.**
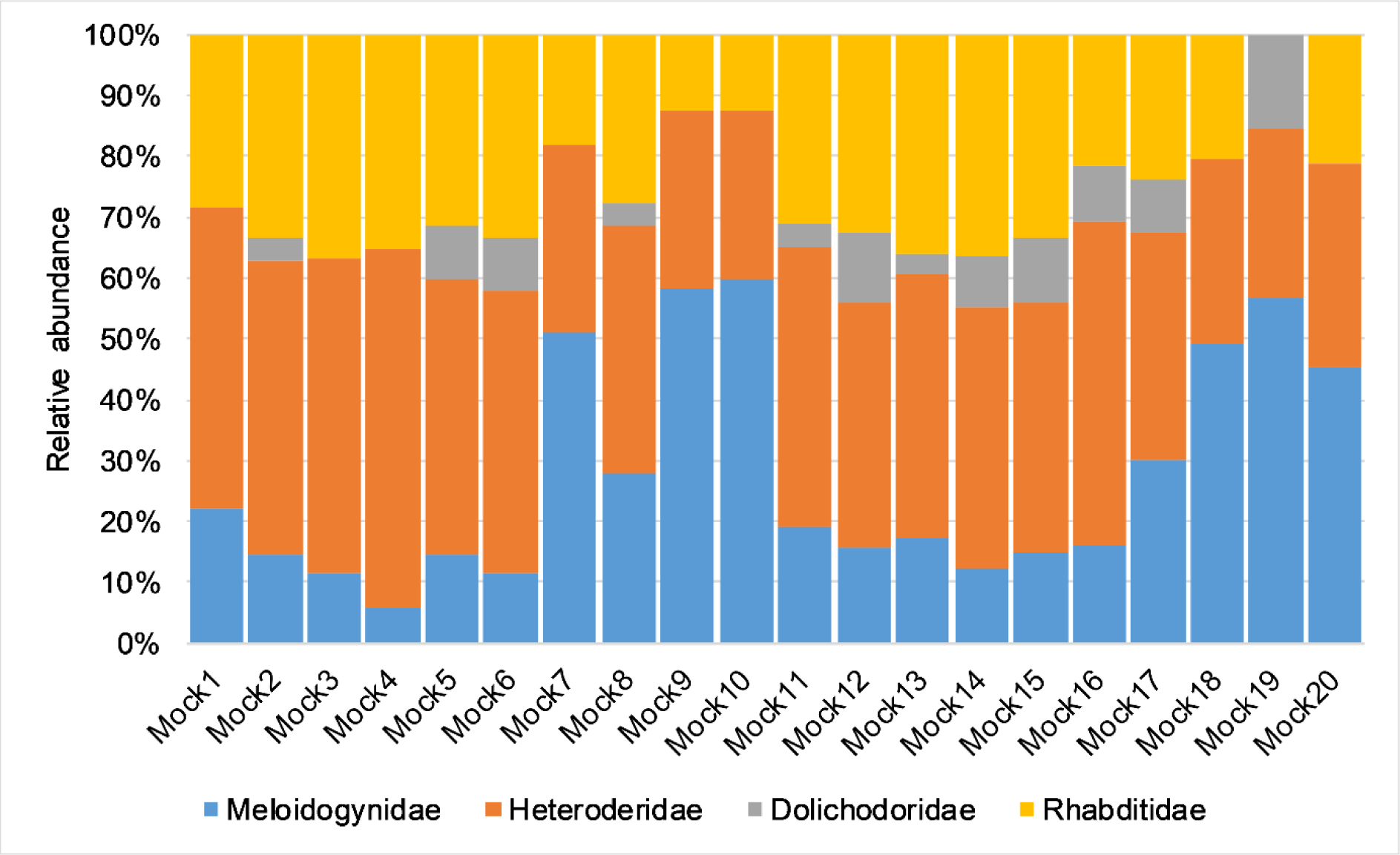
Relative abundance of sequence reads at family rank in mock samples amplified and sequenced using MMS primer set. Only taxonomic assignments appearing in top hits and with sequence similarities ≥ 99% and coverage 100% were considered.

**S3 Fig.**
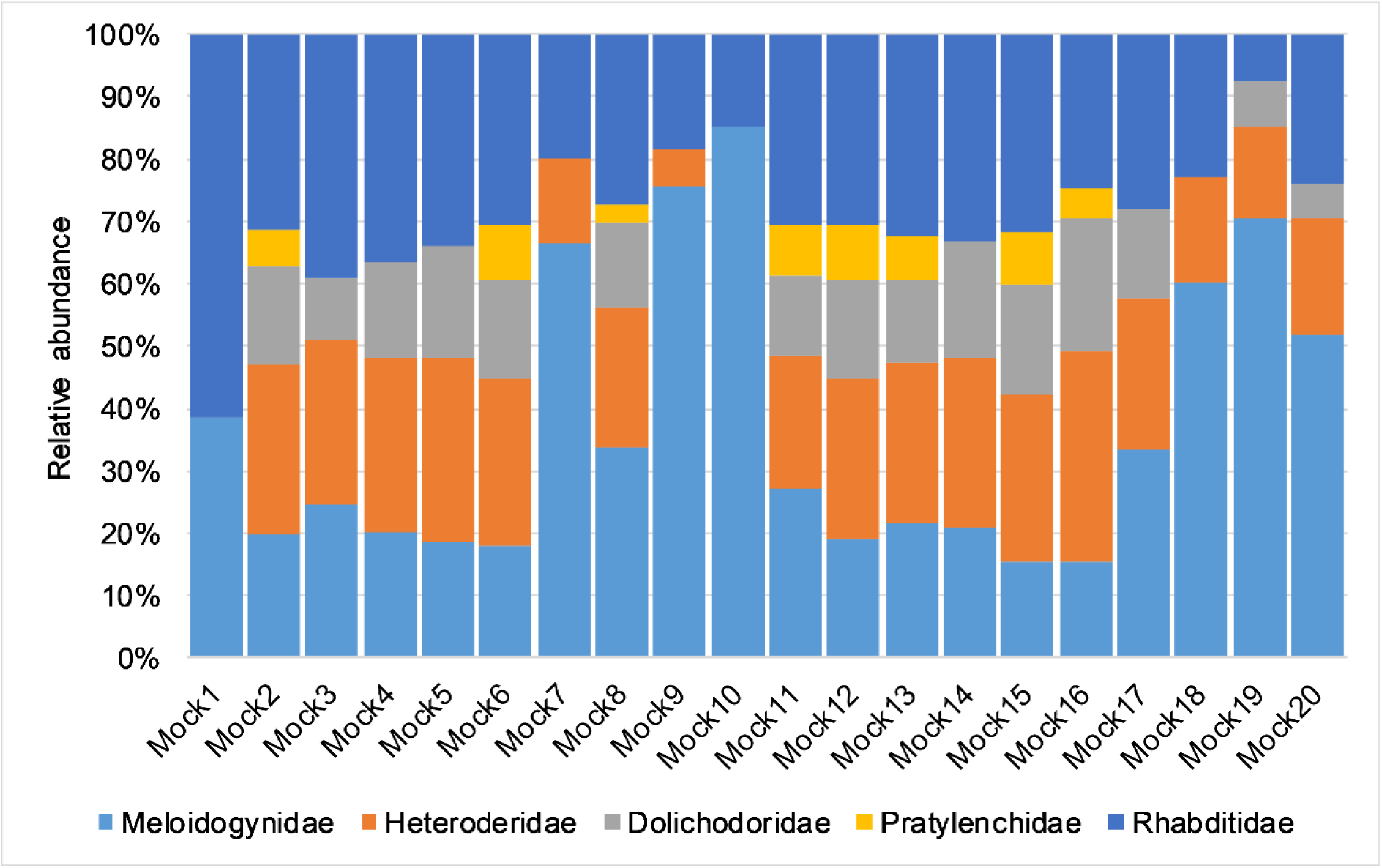
Relative abundance of sequence reads at family rank in mock samples amplified and sequenced using NEM primer set. Only taxonomic assignments appearing in top hits and with sequence similarities ≥ 99% and coverage 100% were considered.

**S4 Fig.**
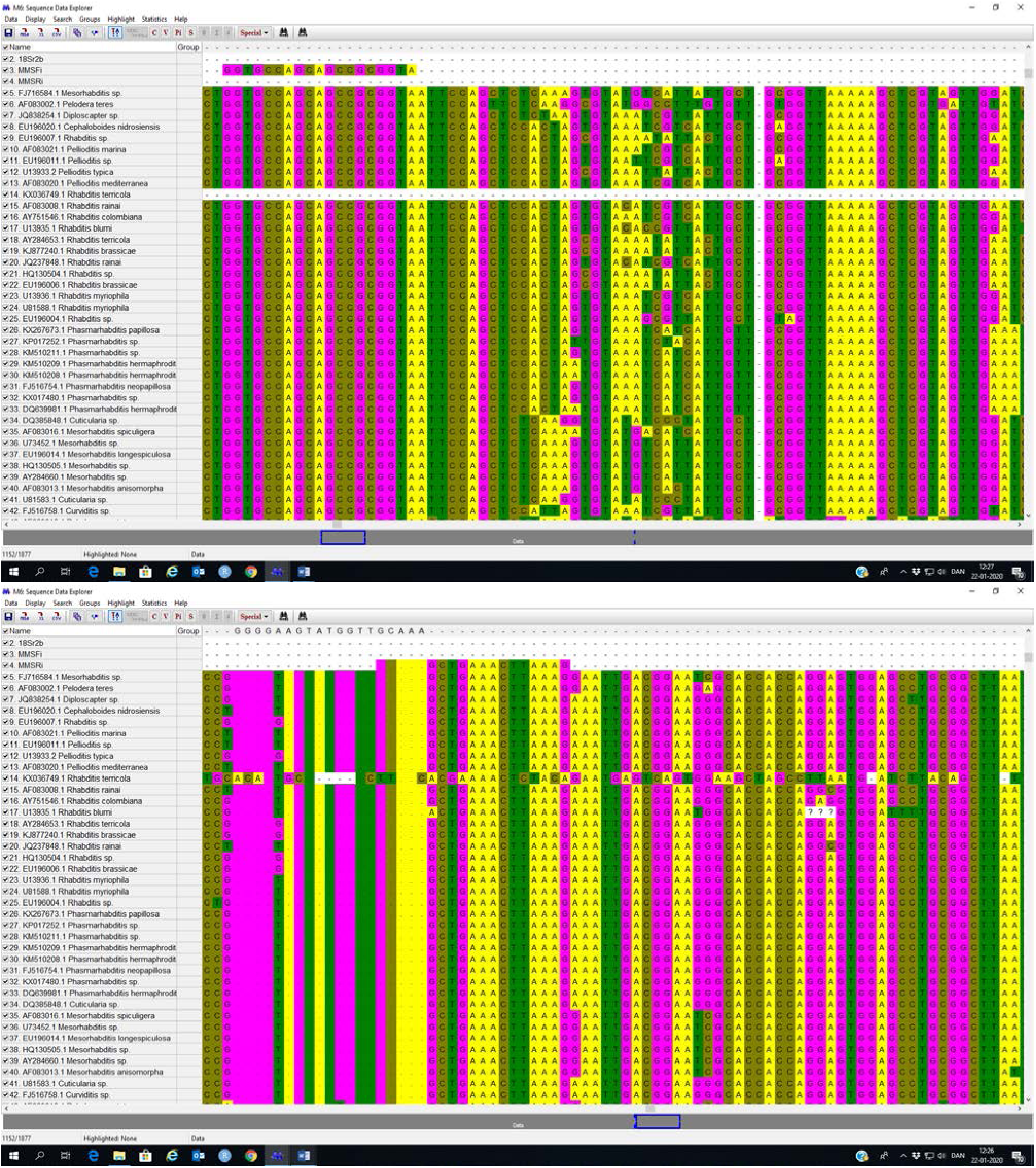
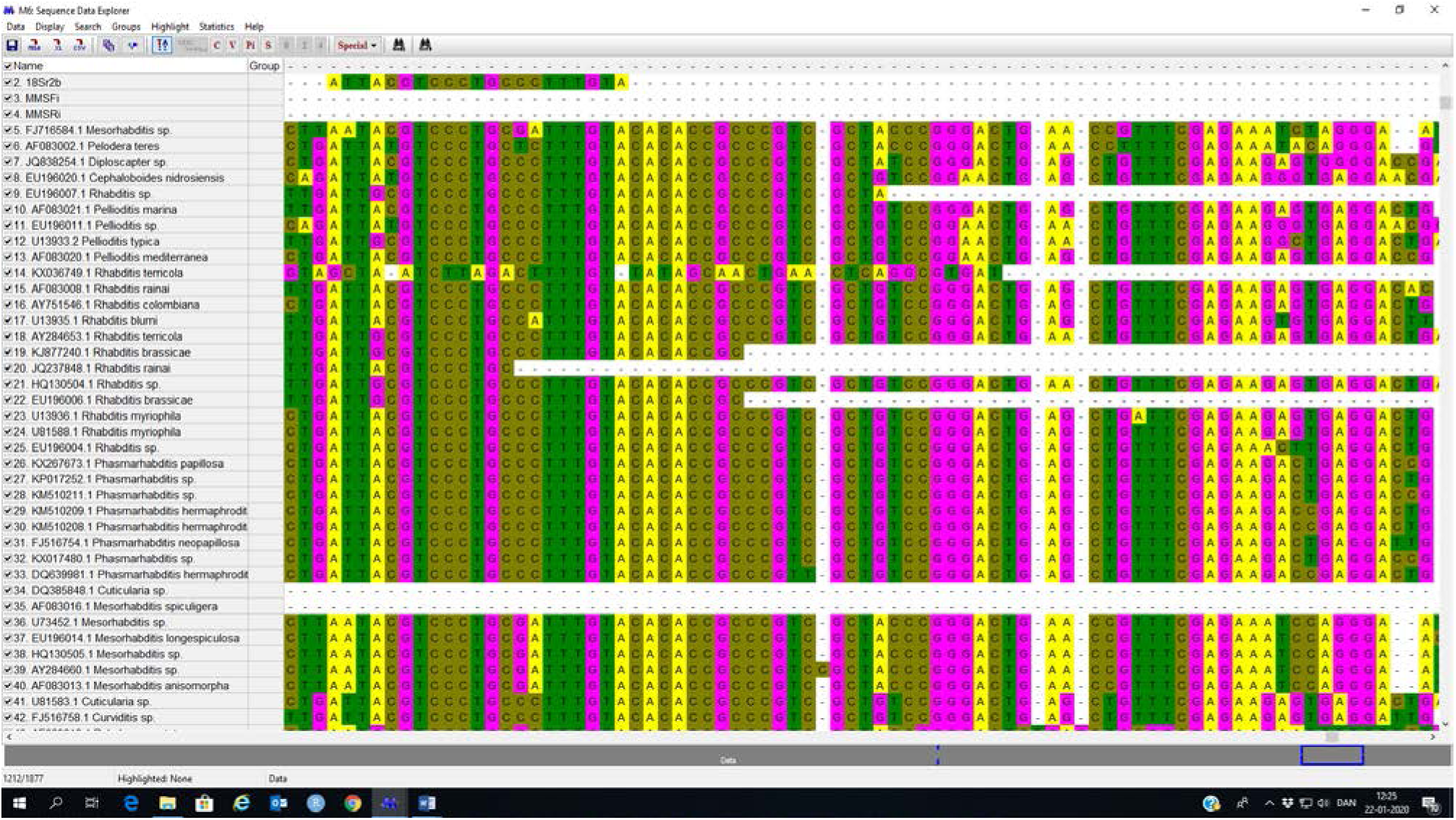
Multiple sequence alignment of MMS and NEM primer sets and representative taxa of Rhabditidae.

